# Reward does not facilitate visual perceptual learning until sleep occurs

**DOI:** 10.1101/738765

**Authors:** Masako Tamaki, Aaron V. Berard, Tyler Barnes-Diana, Jesse Siegel, Takeo Watanabe, Yuka Sasaki

**Affiliations:** Department of Cognitive, Linguistic, and Psychological Sciences, Brown University, 190 Thayer Street, Providence, RI 02912, USA

**Keywords:** Perceptual learning, reward, REM sleep

## Abstract

A growing body of evidence indicates that visual perceptual learning (VPL) is enhanced by reward provided during training. Another line of studies has shown that sleep following training also plays a role in facilitating VPL, an effect known as the offline performance gain of VPL. However, whether the effects of reward and sleep interact on VPL remains unclear. Here, we show that reward interacts with sleep to facilitate offline performance gains of VPL. First, we demonstrated a significantly larger offline performance gain over a 12-h interval including sleep in a reward group than that in a No-reward group. However, the offline performance gains over the 12-h interval without sleep were not significantly different with or without reward during training, indicating a crucial interaction between reward and sleep in VPL. Next, we tested whether neural activations during posttraining sleep were modulated after reward was provided during training. Reward provided during training enhanced REM sleep time, increased oscillatory activities for reward processing in the prefrontal region during REM sleep, and inhibited neural activation in the untrained region in early visual areas in NREM and REM sleep. The offline performance gains were significantly correlated with oscillatory activities of visual processing during NREM sleep and reward processing during REM sleep in the reward group but not in the No-reward group. These results suggest that reward provided during training becomes effective during sleep, with excited reward processing sending inhibitory signals to suppress noise in visual processing, resulting in larger offline performance gains over sleep.

**Significance statement:** Independent lines of research have shown that visual perceptual learning (VPL) is improved by reward or sleep. Here, we show that reward provided during training increased offline performance gains of VPL over sleep. Moreover, during posttraining sleep, reward was associated with longer REM sleep, increased activity in reward processing in the prefrontal region during REM sleep, and decreased activity in the untrained region of early visual areas during NREM and REM sleep. Offline performance gains were correlated with modulated oscillatory activity in reward processing during REM sleep and visual processing during NREM sleep. These results suggest that reward provided during training becomes effective on VPL through the interaction between reward and visual processing during sleep after training.

## INTRODUCTION

Visual perceptual learning (VPL) is defined as the long-term performance improvement on a perceptual task as a result of perceptual experience (1–3). VPL is regarded as a manifestation of plasticity in visual information processing and the brain; however, the neural mechanisms underlying VPL are not completely understood. Recently, the effects of two factors, sleep and reward, on VPL have attracted considerable attention in different contexts, as described below.

First, VPL is considered to have several phases, including the fast within-session phase and the delayed offline phase (4), the latter of which sleep plays an important role (5). Evidence for the delayed offline phase includes the fact that performance gains in VPL emerge overnight (6–8) or after daytime naps (9, 10). Moreover, deprivation of a total night of sleep (11), only REM sleep (6) or only slow wave sleep (stage N3) (12) nullifies the VPL performance gains achieved during sleep. The offline performance gain by sleep has been found in various types of VPL tasks (6–8, 10), as well as other types of learning and memory tasks, including motor skill learning and declarative memory (13–16). Since the mere passage of time, which does not include sleep, shows no clear offline gain of VPL, the offline performance gains of VPL have been proposed to be sleep dependent, not time dependent, and the brain status in sleep itself is believed to be essential for the offline gain of VPL (11, 12, 17).

Another line of research demonstrates that reward provided during training enhances VPL, even without an active task or even when the main visual stimuli were invisible (18–21). For instance, in a previous study (18), as a result of passive viewing of a sequence of two visible or invisible orientations, one paired with reward and the other paired with no reward, only the orientation paired with reward was learned. Because VPL was formed by reward without an active task, Seitz et al. (2009) suggested that the effect of reward on VPL is not due to attention or task-related reinforcement signals but is consistent with a process that gates learning originating in subcortical reward processing or reinforcement learning (22).

Importantly, whether, and if so, how, the effects of reward and sleep interact on VPL has remained unclear because in studies examining the effect of reward on VPL, possible interactions of sleep with reward were not considered and hence not examined. Because VPL is defined as a long-lasting effect, the effect of reward on VPL was reported as a result of multiple-day training periods. For example, the abovementioned study (18) used a training period that covered several days, during which sleep was not experimentally deprived. Therefore, the experiments inevitably include several hours of sleep episodes repeatedly after daily training. Thus, whether these results are likely to reflect interactions between reward and sleep on VPL remains to be tested.

The present study aimed to investigate whether sleep and reward interact on VPL and, if so, how facilitation of VPL is manifested in brain oscillations during sleep. In Experiment 1, we systematically compared the results from 4 conditions in separate groups: a group with neither reward nor sleep, a group with reward without sleep, a group with sleep without reward, and a group with both reward and sleep. We found that reward significantly interacts with sleep, enhancing offline performance gains on a visual task over those achieved by sleep alone but not enhancing performance during training. These results suggest that reward becomes effective during the delayed phase and that sleep facilitates the effect of reward on VPL. In Experiment 2, to test whether the results of Experiment 1 was simply due to reward provided during test sessions in reward groups, reward was provided only during training and not test sessions. We have obtained the same pattern of results as in Experiment 1 in which reward was provided during test sessions as well as training. This indicates that the results of Experiment 1 cannot be simply attributed to reward provided during test sessions. In Experiment 3, we investigated the neural mechanism underlying the interaction between reward and sleep on VPL using polysomnography. Posttraining REM sleep became longer when reward was provided during training than when reward was not provided during training. Additionally, spontaneous brain oscillations that represent reward processing in the prefrontal region (23) and visual processing in early visual areas were modulated in the posttraining sleep. Importantly, modulated spontaneous oscillations for the reward and visual processing were strongly correlated with offline performance gains in the reward group. These results suggest that reward becomes effective through posttraining sleep, during which crucial interactions occur between reward processing and visual processing, which result in enhanced offline performance gains.

## RESULTS

### Experiment 1

We tested whether the effects of reward and sleep on VPL are interactive. Because sleep plays a critical role in offline performance gain, we tested whether offline performance gains were modulated if reward and sleep do in fact interact with each other. By contrast, if the impacts of reward and sleep are independent, then offline performance gains of VPL over sleep should not be significantly different between training with and without reward.

There were 4 groups (Fig. 1; Sleep & Reward, Sleep & No-reward, Wake & Reward, and Wake & No-reward) in which we manipulated sleep and reward factors. We measured performance gains over a 12-h interval. Each group had 2 sessions (1st training and 2nd training), which were separated by a 12-h interval. Two sleep groups (Sleep & Reward and Sleep & No-reward) conducted their 1st training session at 9 pm and their 2nd session at 9 am the following morning, whereas the other 2 awake groups (Wake & Reward and Wake & No-reward) went through the same procedure except that they began their 1st training session at 9 am and the 2nd session at 9 pm on the same day.

**Fig. 1.**
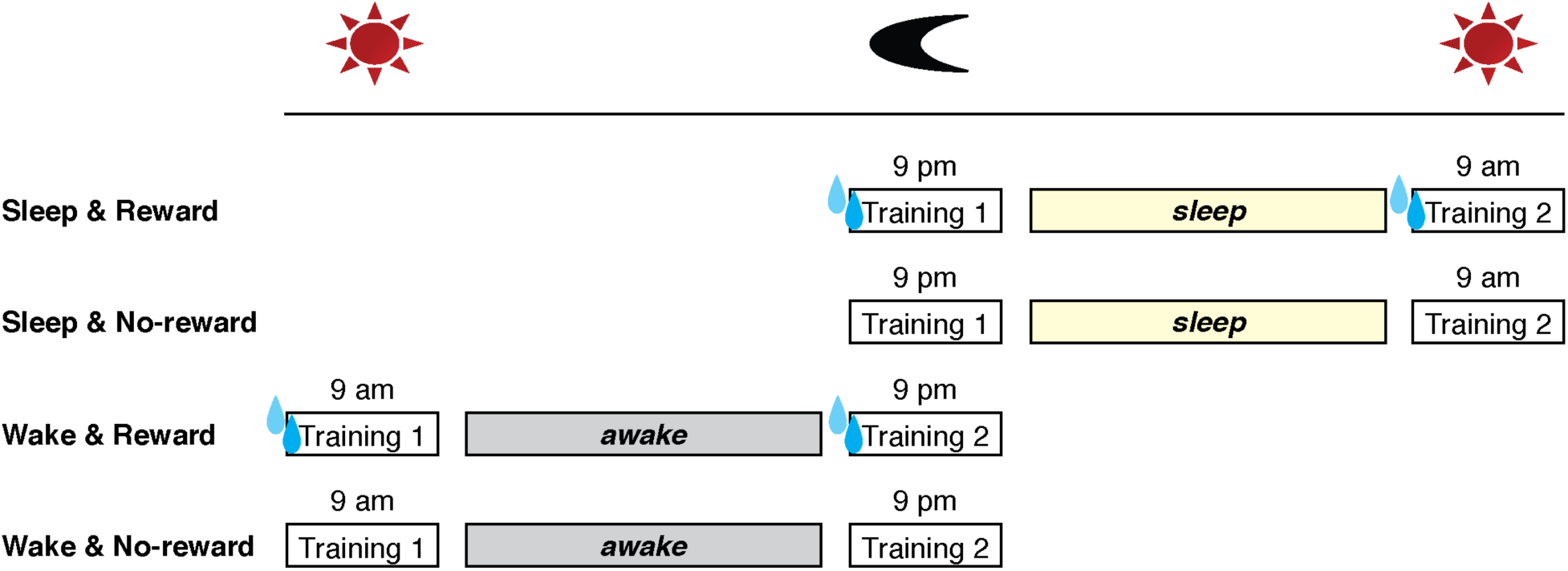
Experimental design for Experiment 1.

For training, we used a texture discrimination task (TDT, see **Materials and Methods**), which is a standard VPL task (6, 9, 12, 24–26). The two reward groups (Sleep & Reward and Wake & Reward) received a water reward upon a correct response in each trial during the 1st and 2nd training sessions (see **Materials and Methods**), whereas the No-reward groups (Sleep & No-reward and Wake & No-reward) did not receive a water reward during training. Subjects in the reward groups were asked to refrain from eating or drinking for 4 h prior to the sessions so that the water reward would be effective.

We conducted a 2-way ANOVA with the factors Sleep (sleep vs. wake) and Reward (reward present or absent) on offline performance gains between the 1st and 2nd training sessions. The mean performance change% (see **Materials and Methods**) for each group is shown in Fig. 2A. The results of the ANOVA indicated a significant interaction between the Sleep and Reward factors (F(1,43)=4.207, p=0.046), as well as significant main effects of Sleep (F(1,43)=22.545, p<0.001) and Reward (F(1,43)=5.207, p=0.028). The results support that the effect of sleep and the effect of reward interact with each other on the offline performance gain.

**Fig. 2.**
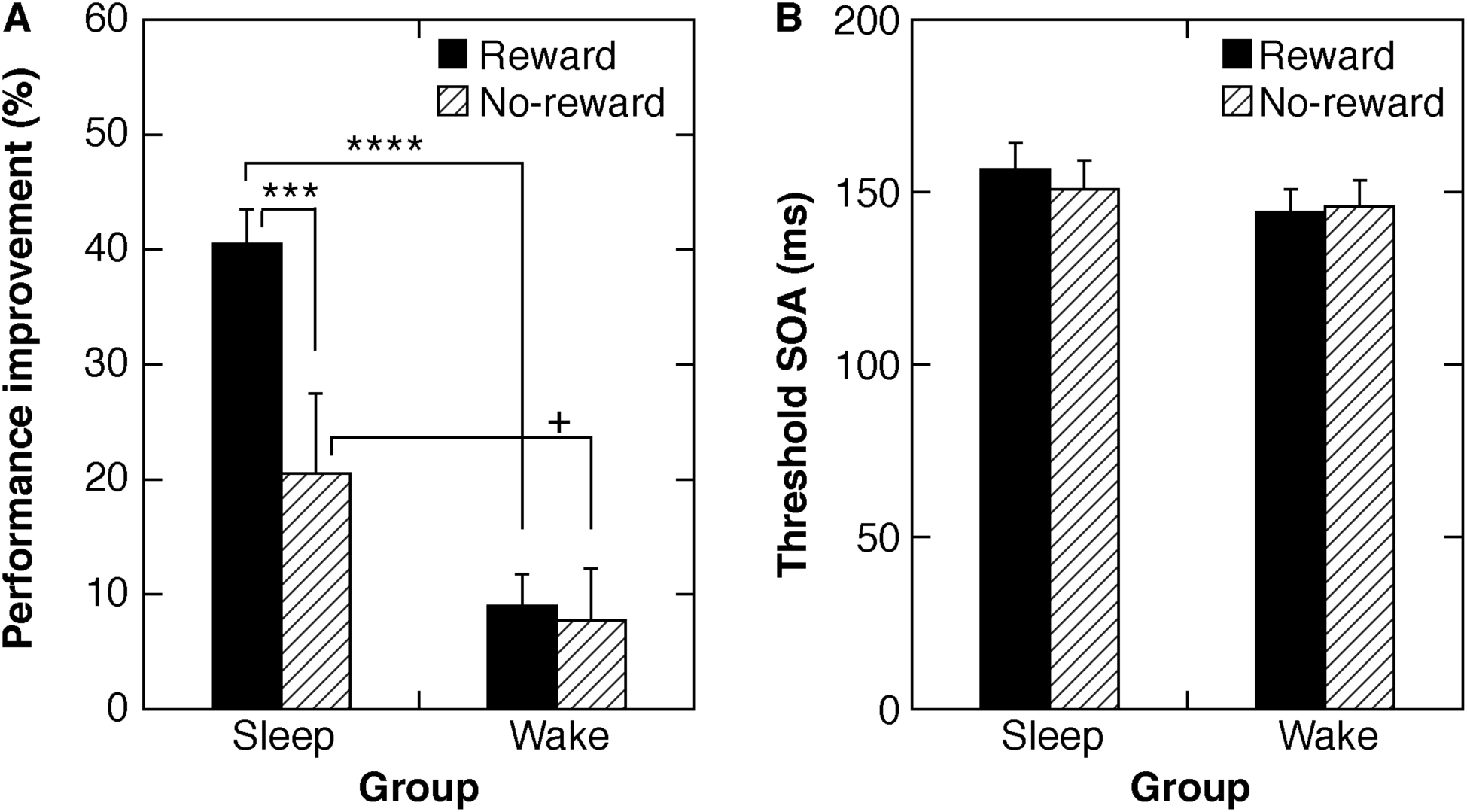
**(A)** Performance improvement (%) in Experiment 1 (mean ± SE). A significant interaction between reward and sleep factors was found (see text). Asterisks indicate statistical results of simple effect tests (**** p<0.001; *** p<0.005; + p<0.1). **(B)** The threshold SOAs (mean ± SE) for the 4 groups in the initial training session were not significantly different.

The results also replicated the effect of sleep on offline performance gains (6–10) and the effect of reward in VPL (18, 19, 21). See **Supplementary Table S1** for the mean threshold SOAs for each training session. Additionally, see **Supplementary Fig. S1** for boxplots of the mean performance improvement%.

To examine the source of the interaction, we further conducted post hoc tests by looking at simple effects. A significant simple effect of Reward was found between the sleep groups (Sleep & Reward vs. Sleep & No-reward; F(1,43)=9.601, p=0.003) but not between the wake groups (Wake & Reward vs. Wake & No-reward; F(1,43)=0.027, p=0.873). These results indicate that the significant interaction between reward and sleep originated from the difference in the reward effect between the sleep and wake groups. Namely, the effect of reward is evident between the sleep groups, while the effect of reward is elusive between the wake groups.

We conducted two types of control analyses. We first tested whether the initial performance, represented by the threshold SOA in the 1st training session, differed across groups. Two factors may have affected the threshold SOAs in the 1st training session. The first is the circadian timing of the sessions, which was different between the sleep and wake groups. The circadian timing might have caused the performance at night to be worse than that during the day (e.g., (27, 28)). The second is that the effect of reward might have accumulated during training and emerged during the 1st training session. If this was the case, the threshold SOA in the 1st training session should be better for the groups who received a reward during training than for those who did not.

We tested whether the threshold SOA during the 1st training (Fig. 2B) was significantly different across conditions via a 2-way ANOVA with factors Sleep and Reward. The results indicated no significant main effect of Sleep (F(1,43)=1.352, p=0.251), no significant main effect of Reward (F(1,43)=0.073, p=0.7881), and no significant interaction of Sleep and Reward (F(1,43)=0.223, p=0.639). These results suggest that neither the difference in the circadian timing nor the reward during training caused performance differences in the 1st training session among groups. The effect of reward did not appear in the first training session but appeared only in the offline performance gains where sleep played a role.

Second, we tested whether the accuracy of the central letter task was the same across groups. If some subjects did not engage well with the central fixation task, that is, they moved their eyes to the orientation task, the performance would be higher during training. We found that the mean accuracy of the central letter task for each group was high during both the 1st and 2nd training sessions. The Shapiro-Wilk tests showed that the distribution of the accuracy of the central task violated normality for the 1st and 2nd training sessions. We thus used the nonparametric Kruskal-Wallis one-way ANOVA to test whether the accuracy of the central task differed across groups for the 1st and 2nd training sessions. The results showed no significant group difference for the 1st (Chi-Square = 3.489, df = 3, p=0.322) or 2nd training sessions (Chi-Square = 1.872, df = 3, p=0.599).

The results of Experiment 1 clearly demonstrate that reward and sleep interact with each other and that the interaction results in larger offline performance gains of VPL. Moreover, the effect of reward is significant for only the sleep group and not for the wake group. These results suggest that the reward may not be effective unless participants sleep after training.

### Experiment 2

In Experiment 1, a reward was provided during both the 1st and 2nd training sessions. Although the reward did not influence the performance during the 1st training session, one may wonder whether the reward given during the 1st and 2nd training sessions accumulated to induce larger performance gains for the groups who received the reward during training. Thus, in Experiment 2, we slightly changed the design. First, we added short test sessions to obtain threshold SOAs separate from a training session. Second, the reward was not given to any of the groups during the test sessions. Thus, the performance evaluation during the test sessions was not influenced by reward.

Two test sessions were performed: a presleep test session (T1) immediately after the training session and a postsleep test session (T2) after a 12-h interval without a second training session (Fig. 3). A reward was not given to any of the groups during the two test sessions.

**Fig. 3.**
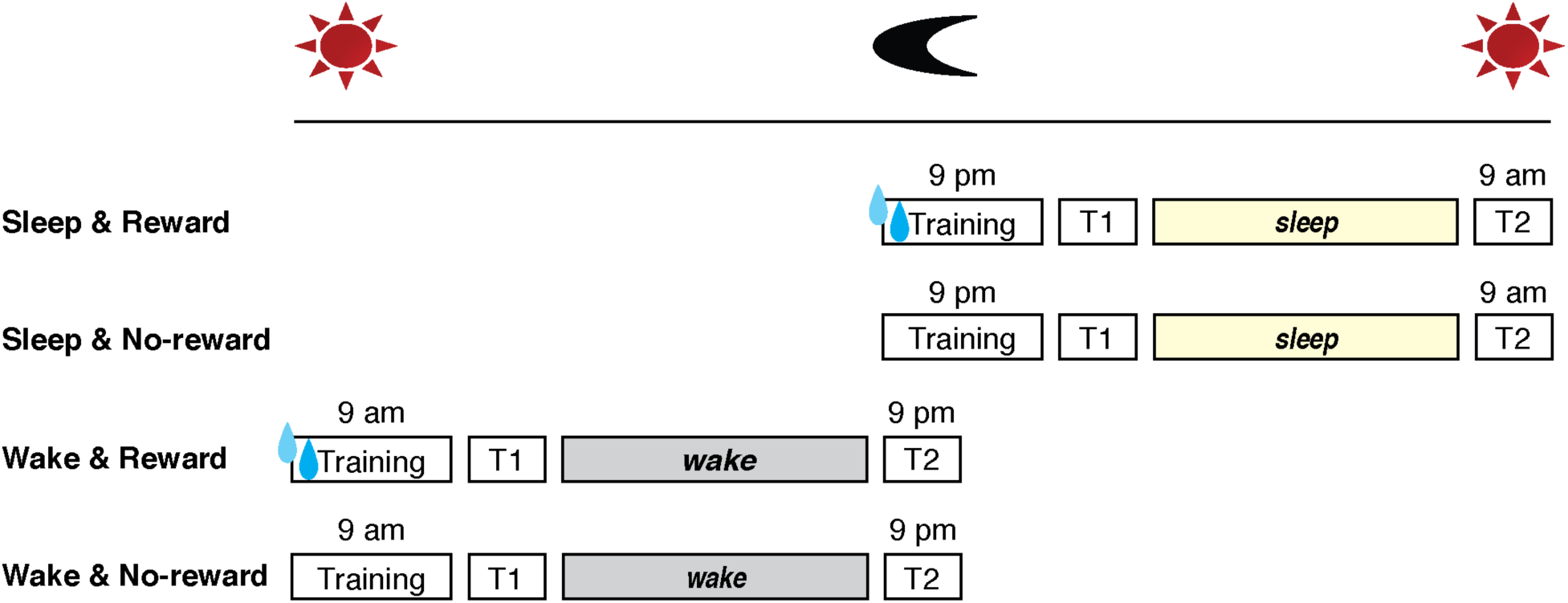
Experimental design for Experiment 2. T1 and T2 represent the first and second test sessions, respectively.

There were 4 groups (Fig. 3; Sleep & Reward, Sleep & No-reward, Wake & Reward, and Wake & No-reward). The two sleep groups (Sleep & Reward and Sleep & No-reward) performed their training at 9 pm followed by test session T1 and test session T2 at 9 am the following morning, whereas the other two wake groups (Wake & Reward and Wake & No-reward) followed the same procedure except that they performed their training at 9 am in the morning followed by test session T1 and then performed test session T2 at 9 pm later the same day. The procedures for the VPL task and reward were the same as those in Experiment 1.

We conducted a 2-way ANOVA with factors Sleep (sleep vs. wake) and Reward (present or absent) on the offline performance gains between test sessions T1 and T2. See **Supplementary Table S2** for the mean threshold SOA for the test and training sessions. The mean offline performance gains for each group are shown in Fig. 4A and **Supplementary Fig. S2**. The results of the ANOVA with Sleep and Reward factors replicated the significant interaction between Sleep and Reward on offline performance gains (F(1,36)=6.041, p=0.019), as well as a significant main effect of Sleep (F(1,36)=37.321, p<0.001). A main effect of Reward was not significant (F(1,36)=0.257, p=0.615). In addition, a significant simple effect of Reward was found between the sleep groups (Sleep & Reward vs. Sleep & No-reward, F(1,20)=7.100, p=0.015) but not between the wake groups (Wake & Reward vs. Wake & No-reward, F(1,16)=1.238, p=0.282). These findings replicated the interaction between reward and sleep in Experiment 1, showing that reward has no effect on the offline performance gains of VPL without sleep.

**Fig. 4.**
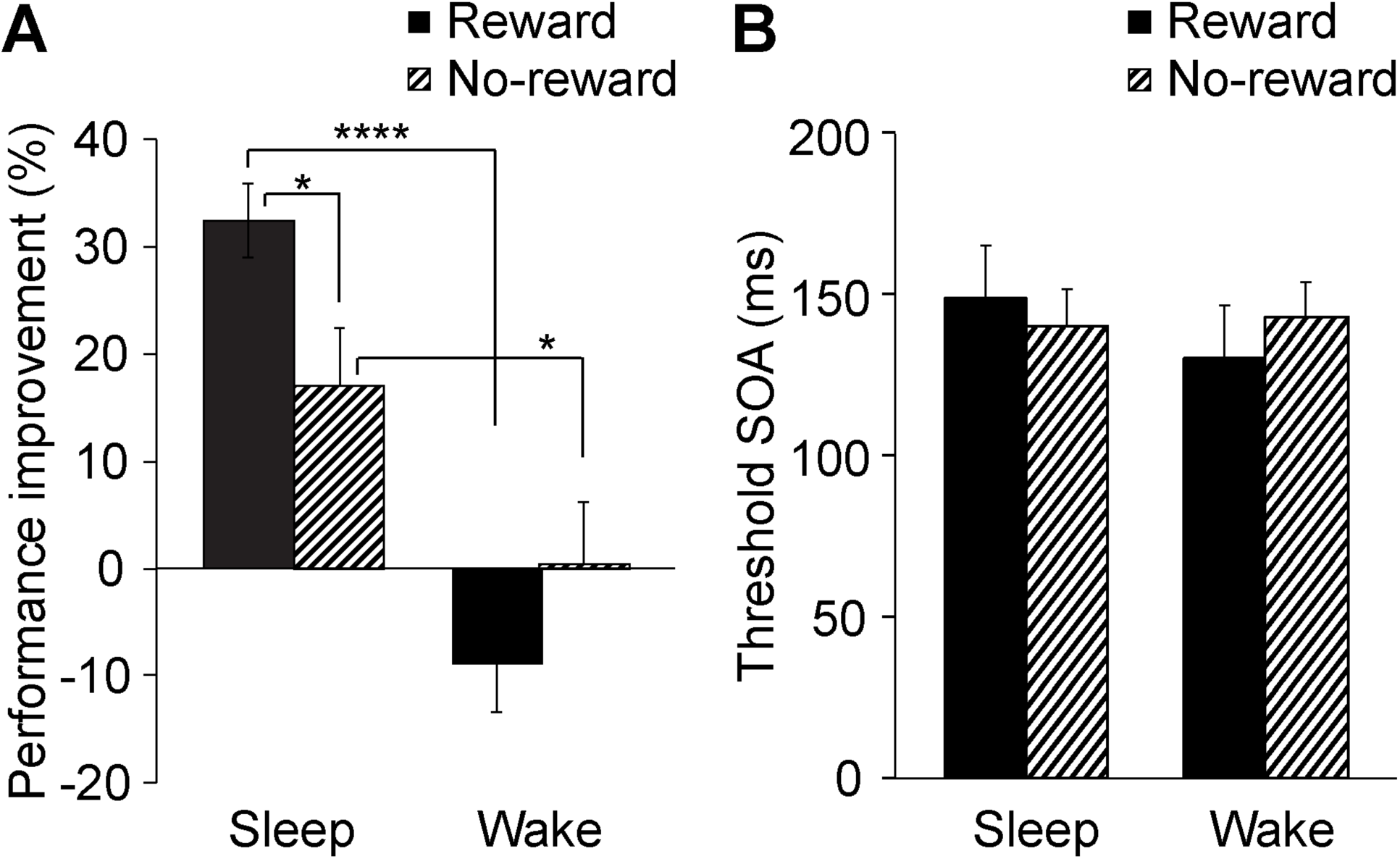
**(A)** Performance improvement (%) in Experiment 2 (mean ± SE). A significant interaction between reward and sleep factors was found (see text). Asterisks indicate statistical results of simple effect tests (****p<0.001, * p<0.05). **(B)** The threshold SOAs (mean ± SE) for the 4 groups in the initial training session were not significantly different.

We also conducted control analyses. First, we tested whether the performance in the initial training session differed across groups. Fig. 4B plots the threshold SOA for the initial training session for all 4 groups. A 2-way ANOVA with factors Sleep and Reward applied to the threshold SOA indicated no significant main effect of Sleep (F(1,36)=0.049, p=0.827), no significant main effect of Reward (F(1,36)=0.267, p=0.608), and no significant interaction of Sleep and Reward (F(1,36)=0.030, p=0.864). In addition, the threshold SOA during test session T1 was not significantly different across the 4 groups. A 2-way ANOVA with factors Sleep and Reward applied to the threshold SOA during test session T1 indicated no significant main effect of Sleep (F(1,36)=0.164, p=0.688), no significant main effect of Reward (F(1,36)<0.001, p=0.994), and no significant interaction of Sleep and Reward (F(1,36)=0.881, p=0.354). These analyses indicate that the performance was not significantly different across groups during the training session and test session T1.

Second, the central task performance was not significantly different across groups after training. Again, since the Shapiro-Wilk tests showed that the data violated normality, we used a Kruskal-Wallis one-way ANOVA to test whether the accuracy of the central task differed across groups for the 1st and 2nd training session. The results showed no significant group difference for the 1st (Chi-Square = 0.7, df = 3, p=0.873) or 2nd training sessions (Chi-Square = 0.846, df = 3, p=0.838).

Notably, the number of trials used to obtain the threshold SOAs was 10 in Experiment 2, which was smaller than the 78 trials used in Experiment 1. Some might wonder whether the smaller number of trials caused too much variability and lowered the precision of the measurement. We examined whether different numbers of trials resulted in different threshold SOAs in Experiment 2. We computed the threshold SOAs using the first 10 trials as well as using 78 trials in the training session in Experiment 2. We then used one-way repeated measures ANOVA (factor = # of trials) to test whether the threshold SOAs measured using different numbers of trials were significantly different. No significant difference was observed between the threshold SOAs based on 10 trials vs. 78 trials (F(1,39)=0.142, p=0.708). In addition, the variability for the threshold SOA was not significantly different between the 10 trials and 78 trials (Barlett’s statistic = 0.343, p = 0.558). Therefore, using a smaller number of trials did not cause a serious problem with obtaining the threshold in Experiment 2.

### Experiment 3

The findings thus far led to a hypothesis that the interaction between reward processing and visual processing takes place during posttraining sleep after reward is given during training. In Experiment 3, we examined whether this is the case, and if so, how the interaction occurs in the brain during sleep using polysomnography (PSG, see **Materials and Methods**).

We examined whether spontaneous brain oscillations in reward and visual processing during posttraining sleep (a nap) were modulated by reward. Since we were interested in the interaction between reward and sleep, only two sleep groups were considered: with and without reward during the training session (Fig. 5; Reward vs. No-reward). The design in Experiment 3 was similar to that in Experiment 2. The presleep test session (T1) was conducted immediately after the training session. The postsleep test session (T2) was conducted after the nap (Fig. 5). No reward was given during the test sessions.

**Fig. 5.**
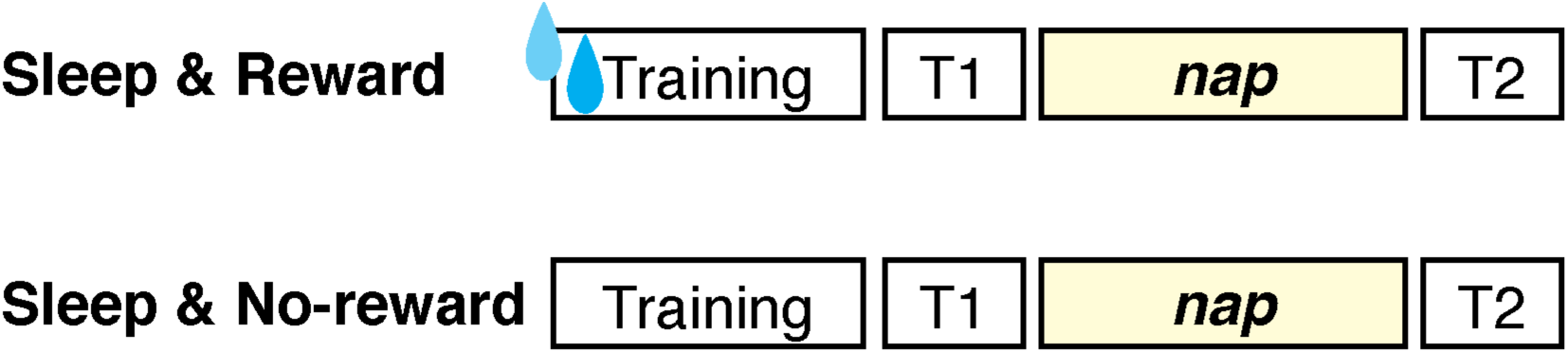
Experimental design for Experiment 3. T1 and T2 represent the first and second test sessions, respectively.

### Sleepiness between groups

We measured subjective sleepiness by the Stanford Subjective Sleepiness (SSS) scale (29) at the beginning of sessions T1 and T2. See **Supplementary Table S3** for the SSS results. We tested whether sleepiness differed between groups. The subjective sleepiness was not significantly different between the groups at T1 (Wilcoxon-Mann-Whitney test, p=0.852) or T2 (Wilcoxon-Mann-Whitney test, p=0.775). Thus, differential behavioral outcomes or spontaneous oscillations between groups could not be attributed to sleepiness.

### Performance gain

Both groups showed significant offline performance gains over the sleep period (Fig. 6; one-sample t-test; Reward group: t(10)=8.22, p<0.001; No-reward group: t(10)=3.13, p=0.011). One-way ANOVA with the factor being Reward (reward present vs. absent) on the offline performance gain showed that the main effect of Reward was significant (F(1,20)=4.655, p=0.043), replicating the previous finding that the offline performance gain was larger for the Reward group than for the No-reward group (Fig. 6A; see **Supplementary Fig. S3** for boxplots and **Supplementary Table S4** for the threshold SOA (ms) for the training, 1st and 2nd test sessions).

**Fig. 6.**
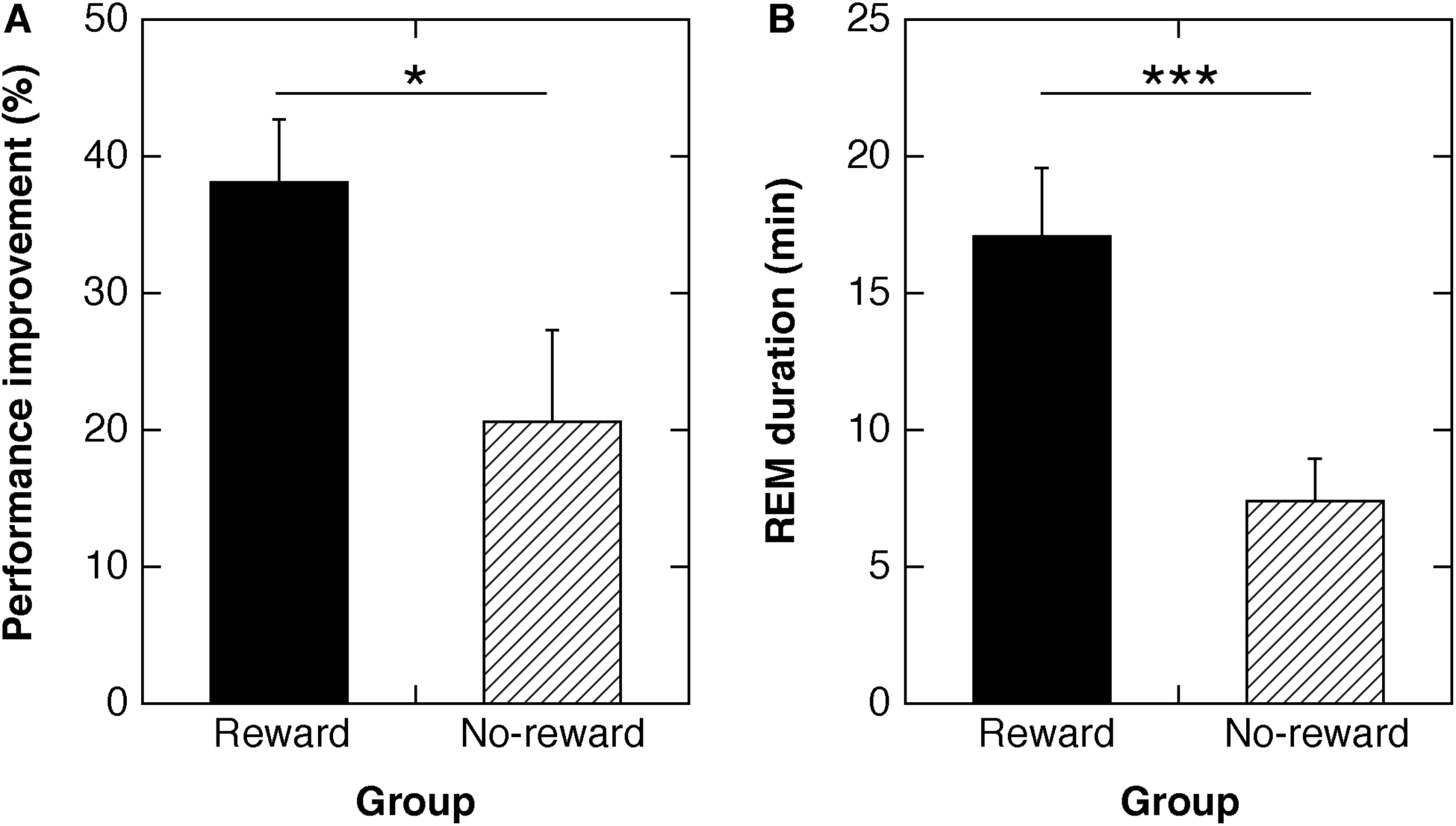
**(A)** Performance improvement% in Experiment 3 (mean ± SE). * p<0.05. **(B)** REM sleep duration (min) (mean ± SE). *** p<0.005.

### Time spent in each sleep stage

We investigated whether the sleep structures, which included 7 variables (see Table 1, time spent (min) in Wake, N1, N2, N3, and REM sleep, sleep-onset latency (SOL) and REM sleep latency in the posttraining nap, see **Materials and Methods**), were different between the Reward and No-reward groups. To do this analysis, we repeated the one-way ANOVA 6 times. Only the time spent in REM sleep was significantly different between groups (Fig. 6B, Table 1).

**Table 1.** Sleep structures (min) for Reward and No-reward groups

|  | Reward |  |  | No-reward |  |  |  |
| --- | --- | --- | --- | --- | --- | --- | --- |
| SOL | 9.96 | ± | 1.59 | 8.73 | ± | 1.59 | t(20)=0.542, p=0.594 |
| Wake | 11.55 | ± | 2.57 | 10.05 | ± | 3.11 | t(20)=0.372, p=0.714 |
| N1 | 9.23 | ± | 1.54 | 8.73 | ± | 1.63 | t(20)=0.223, p=0.826 |
| N2 | 31.05 | ± | 2.09 | 36.86 | ± | 5.00 | t(20)=1.073, p=0.296 |
| N3 | 20.86 | ± | 3.17 | 23.73 | ± | 4.57 | t(20)=0.515, p=0.612 |
| REM | 16.95 | ± | 2.52 | 7.32 | ± | 1.59 | t(20)=3.231, p=0.004 |
| REM latency<sup>a</sup> | 62.41 | ± | 6.05 | 68.44 | ± | 3.36 | t(18)=0.818, p=0.424 |

Values are the mean ± SE. SOL: sleep-onset latency. <sup>a</sup>REM latency: Latency to the onset of REM sleep. The number of subjects for the REM latency was n=11 for the Reward group and n=9 for the No-reward group since 2 subjects in the No-reward group did not show REM sleep.

### Spontaneous oscillations in the prefrontal and occipital cortex

Next, we investigated whether the activations of spontaneous oscillations during posttraining sleep were different between the Reward and No-reward groups. If reward provided during training induces interactions between reward processing and visual processing during sleep, the interaction would be shown in differential brain activations between the Reward and No-reward groups, depending on sleep stages or frequency bands.

We obtained the power densities for the 4 frequency bands (power densities for delta, theta, alpha, and sigma, see **Materials and Methods**) of spontaneous oscillations from 3 brain regions (prefrontal region and trained and untrained occipital regions) for both NREM sleep and REM sleep. We preselected the prefrontal region because we were interested in reward processing, which is reflected by the strength of spontaneous oscillations in the prefrontal EEG channels (23). Additionally, we preselected the occipital region for visual processing. We used the trained and untrained hemispheres of the occipital EEG channels corresponding to early visual areas, which are known to be involved in the performance gains of this task (30, 31) (see **Materials and Methods**).

To analyze whether reward provided during training modulates spontaneous oscillatory activity in reward and visual processing during posttraining sleep, we conducted a 4-way repeated measures ANOVA with Stage (NREM sleep vs. REM sleep), Frequency (delta, theta, alpha, and sigma bands), Region (prefrontal, trained occipital, and untrained occipital), and Reward (Reward vs. No-reward groups) factors (Fig. 7) on power density. Importantly, the 4-way interaction was significant (F(6,108)=3.234, p=0.006). The significant 4-way interaction allowed us to perform further *post hoc* analyses without inflating the type I error rates. See **Supplementary Table S5** for the complete results of the four-way and post hoc one-way ANOVAs, including 3-way interactions, 2-way interactions and main effects.

**Fig. 7.**
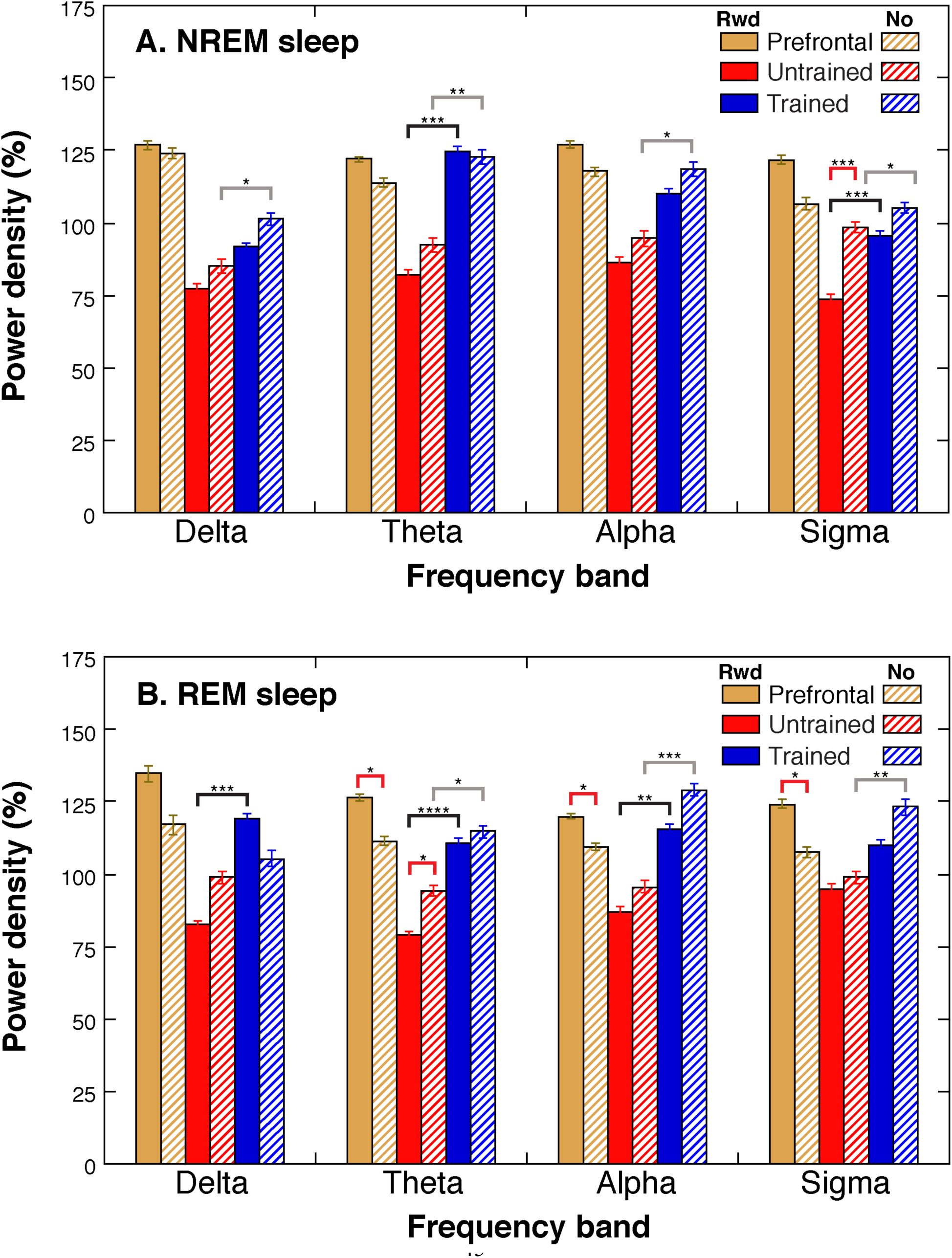
Spontaneous activities (mean ± SE) during NREM sleep **(A)** and REM sleep **(B)**. Filled bars show the Reward group, and hatched bars show the No-reward group. Yellow bars represent prefrontal, red bars represent untrained occipital, and blue bars represent trained occipital regions. A red bracket indicates a significant reward effect in the post hoc analyses. A black bracket shows a significant difference in power densities between the trained and untrained occipital regions in the Reward group, while a gray bracket shows a significant difference in power densities between the trained and untrained occipital regions in the No-reward group in the post hoc analyses. Asterisks indicate statistical significance in the post hoc analysis. See text for details of the statistics. * p<0.05, ** p<0.01, *** p<0.005.

Since we were interested in the reward effect, we conducted one-way ANOVAs to test whether the power density of each frequency band for each sleep stage for each ROI was different between the Reward and No-reward groups as post hoc tests (see **Supplementary Table S5B** for more details). We found that during NREM sleep, only the sigma band of the untrained occipital region showed a significant main effect of Reward (F(1,18)=12.434, p=0.002). During REM sleep, the effect of Reward was observed not only for the untrained occipital region but also for the prefrontal region. A significant main effect of Reward was found at the prefrontal region for the theta (F(1,18)=5.423, p=0.032), alpha (F(1,18)=5.256, p=0.034) and sigma (F(1,18)=4.718, p=0.043) bands, as well as at the untrained occipital region for the theta (F(1,18)=4.822, p=0.041) band during REM sleep.

Notably, the effect of reward on spontaneous oscillations differed for the prefrontal and occipital regions. The power densities of the prefrontal region tended to be larger in the Reward group than in the No-reward group, while the power densities of the occipital regions in the Reward group were smaller than those in the No-reward group (Fig. 7). Because the 4-way interaction was significant and the 2-way interaction between Region and Reward was almost significant (**Supplementary Table S5A**), we tested the effect of reward on the averaged power density across the 4 bands at the prefrontal region and at occipital regions separately for NREM and REM sleep as post hoc tests (**Supplementary Fig. S4**).

For the prefrontal region, a one-way ANOVA with a Reward factor (Reward vs. No-reward) on the averaged power density showed a tendency of a main effect of Reward during REM sleep (F(1,18)=4.358, p=0.051, **Supplementary Fig. S4B**) but not during NREM sleep (F(1,18)=1.872, p=0.188, **Supplementary Fig. S4A**).

For the occipital regions, a two-way ANOVA with Region (Trained vs Untrained) and Reward (Reward vs No-reward groups) factors on the averaged power density showed a tendency of a main effect of Reward during NREM sleep (F(1,18)=3.455, p=0.0795, **Supplementary Fig. S4C**) but not during REM sleep (F(1,18)=1.519, p=0.2336, **Supplementary Fig. S4D**). These results suggest that although not statistically significant, the effect of reward tended to be excitatory in the prefrontal region and was more apparent during REM sleep, whereas the effect of reward tended to be inhibitory in the occipital regions, which was more apparent during NREM sleep.

Previous studies showed a trained-region-specific effect of VPL in early visual areas during sleep (30, 32). We also replicated the effect. Notably, the abovementioned 2-way ANOVA on the averaged power density at the occipital regions showed a significant main effect of region during NREM sleep (F(1,18)=16.752, p=0.0007) and during REM sleep (F(1,18)=25.659, p=0.0001). We further averaged the power density across NREM sleep and REM sleep to test whether the averaged power density during both NREM sleep and REM sleep differed significantly between the trained and untrained regions. The averaged power density was significantly different between the trained and untrained occipital regions in both the Reward group (t(10)=3.416, p=0.007) and the No-reward group (t(8)=4.072, p=0.004). These results replicate the trained-region-specific activity during sleep (30, 32).

We tested whether the modulated power density in the prefrontal and occipital regions during posttraining sleep was associated with the offline performance gains in the Reward group. We obtained the power density for reward processing by averaging across the 4 bands in the prefrontal region during NREM and REM sleep in the Reward group. For the visual processing, we calculated the trained-region-specific activation as the averaged power density across all 4 bands in the trained region of the occipital, from which the averaged power density in the untrained regions of the occipital was subtracted for NREM and REM sleep in the Reward group.

We found that the offline performance gains were significantly correlated with the averaged power density at the prefrontal region during REM sleep (r=0.850, n=11, p=0.001, uncorrected for the multiple comparisons; Fig. 8B) and with the trained-region-specific activation during NREM sleep (r=0.804, n=11, p=0.003, uncorrected for the multiple comparisons; Fig. 8A). By contrast, the offline performance gains were not significantly correlated with the averaged activation at the prefrontal region during NREM sleep (r=0.502, n=11, p=0.116) or with the trained-region-specific activation during REM sleep (r=0.494, n=11, p=0.122).

**Fig. 8.**
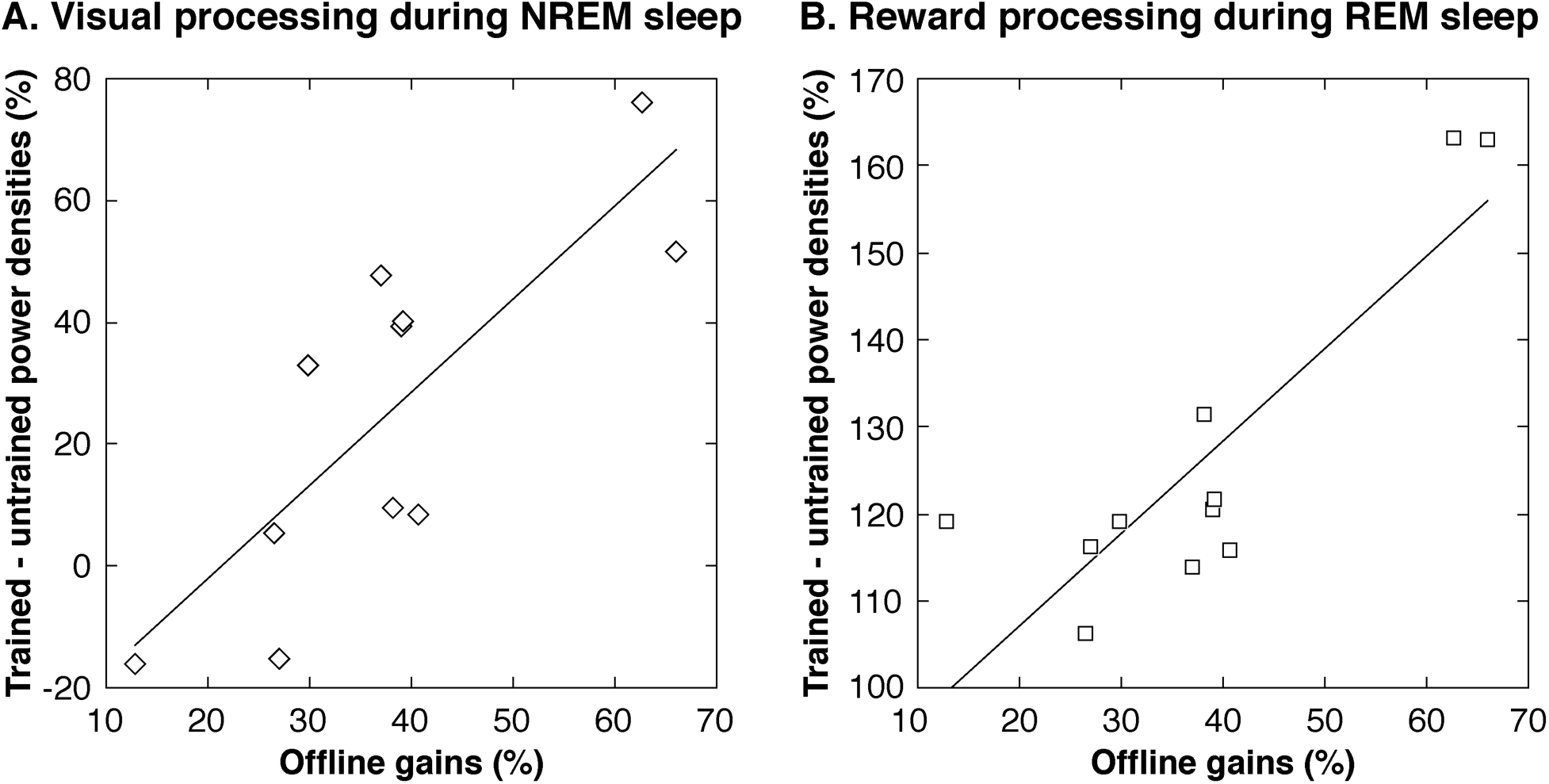
Correlation between offline performance gains and spontaneous oscillations during posttraining sleep in the Reward group (n=11). (**A)** Trained-location-specific power densities in the visual processing during NREM sleep (r=0.804, p=0.003). No outliers were detected by Grubb’s test. (**B)** The prefrontal power densities during REM sleep (r=0.850, p=0.001). No outliers were detected by Grubb’s test.

If these significant correlations are specific to the effect of reward, similar correlations between the oscillatory activations and the offline performance gains should not be seen in the No-reward group. Indeed, the offline performance gains were not significantly correlated with the averaged activation at the prefrontal region during REM sleep (r=-0.348, n=9, p=0.359) or with the trained-region-specific activation during NREM sleep in the No-reward group (r=0.128, n=9, p=0.742).

### Control tests

First, we tested whether the performances during training and at T1 were different between the Reward and No-reward groups. The results of a one-way ANOVA with the factor being Reward (presence vs. absence) on the threshold SOA were not significantly different between the groups during training (F(1,20)=0.310, p=0.912) or during T1 (F(1,20)=0.092, p=0.764).

Second, the central task performance was not significantly different between groups during the test sessions. We analyzed the correct response rates for the central task for all subjects in all groups. Since the Shapiro-Wilk tests showed a violation of normality, we used the Mann-Whitney U test. The results showed no significant difference in the central task performance between the groups in the T1 (U=57, p=0.816) and T2 (U=34, p=0.078) test sessions.

Third, we tested whether any presleep behavioral measures such as the threshold SOA during the training and T1 test sessions were correlated with sleep structures. None of stage W, N1, N2, N3, REM, or sleep-onset latency was significantly correlated with the threshold SOA during training or T1 (see **Supplementary Table S6**).

Finally, we tested whether various sleep habit measures were significantly different between the Reward and No-reward groups. We used the Pittsburg Sleep Quality Index (PSQI) (33) and the Munich Chronotype Questionnaire (MCTQ) (34, 35) to investigate individual habitual sleep quality, habitual bed times, wake-up time, and chronotype. None was significantly different between groups (see **Supplementary Table S7**).

These control analyses suggest that the significant difference in the offline performance gains, as well as the significant differences in the prefrontal and occipital regions between the groups in Experiment 3, were not attributed to performance before sleep, the central task performance, or habitual sleep measures.

## DISCUSSION

The present study clearly demonstrates that an interaction of reward and sleep occurs on VPL: reward provided during training enhanced the offline performance gains achieved by sleep. This result was replicated in three independent experiments. Moreover, the activations of reward processing and visual processing were modulated in posttraining sleep after training with reward.

The effect of reward was two-fold during posttraining sleep: excitation of reward processing in the prefrontal region and inhibition of visual processing, especially in the untrained region of the visual areas. Modulated visual processing during NREM sleep and reward processing during REM sleep were closely correlated with offline performance gains. Note that Experiment 3 used a daytime nap for sleep, while Experiments 1 and 2 used nocturnal sleep. Thus, the interaction between reward and sleep occurs both for daytime naps and nocturnal sleep.

The inhibitory effect of the interaction between reward and sleep on early visual areas was consistent with previous studies. First, prefrontal activation has been shown to provide inhibitory input to sensory cortices (36, 37). Second, reward circuits have been found to send out inhibitory signals (38–40). Third, reward signals selectively decreased blood oxygen level-dependent activation in the early visual areas in a study that used functional magnetic resonance imaging (41). The inhibitory signals sent from reward processing may reduce neural activation in visual processing so that the most efficient neural circuits or synaptic connections in the visual cortex would be preserved, while the neural circuits that are not highly efficient would be trimmed.

In particular, an inhibitory reward effect on the untrained occipital region was found in the sigma band during NREM sleep and in the theta band during REM sleep. These two types of spontaneous oscillations have been linked to learning and memory facilitation during sleep in previous studies. First, sigma activity during NREM sleep, which reflects the activity of sleep spindles, has been linked to neuronal replay or reactivation (42, 43). Sigma activity in the occipital areas is also suggested to be associated with reactivation of neurons in the early visual areas after visual training (32) and in motor areas in posttraining sleep (16, 44), leading to offline performance gains. Second, theta activity during REM sleep has been reported to causally involve contextual memory consolidation in rodents (39) to reorganize firing rates and synchrony of neurons and has been suggested to play a prominent role in REM sleep in sleep-related neuronal plasticity (45). Thus, these two types of spontaneous oscillations, sigma band during NREM sleep and theta band during REM sleep, may be crucially involved in the offline performance gains of VPL over sleep so that the reward further facilitated the roles of these oscillations.

The effect of reward was observed during both NREM sleep and REM sleep. However, the spatiotemporal patterns of modulation by reward may not be the same between NREM and REM sleep. During NREM sleep, the effect of reward was found only in sigma activity in the untrained occipital location. On the other hand, during REM sleep, a wider range of frequency bands, including theta, alpha, and sigma activity, was modulated at the prefrontal region and the untrained region of the occipital cortex after training with reward. This result suggests that the impact of reward may be spatially more diffusive and temporally broader during REM sleep than during NREM sleep to facilitate cortical reorganization and optimization of cortical connectivity to strengthen learning (46–48).

Why did the REM sleep duration increase after training with reward? In the present study, reward provided during training may have excited the neural circuits involved in reward processing. Given that neurons used in prior wakefulness are likely to be reactivated in subsequent sleep (49–52), the excited reward processing during training may have been reactivated during posttraining sleep. Importantly, the neural circuits involved in reward processing largely overlap with the limbic system, which is known to be more active during REM sleep than during NREM sleep (53–57). In addition, two major populations of neurons associated with the onset of REM sleep reside in the posterior hypothalamus (57, 58), which is part of the limbic system. Excitation of the limbic system may provide a brain status that is preferable for maintenance of REM sleep (57), which might result in longer REM sleep duration in the Reward group.

We propose a possible neural mechanism for the effect of reward provided during training to enhance offline performance gains over sleep based on the present results. The offline performance gains would increase because a neural circuit in early visual areas associated with a successful trial during training was tagged by reward during training and would be reactivated during posttraining sleep more than a neural circuit associated with an unsuccessful trial. Reward provided during training would also reactivate reward processing and the limbic system (59) during posttraining sleep. The enhanced reward processing during posttraining sleep would then send inhibitory signals to visual processing to shape and store efficient neural circuits (39, 60) during posttraining sleep, leading to larger offline performance gains by sleep. As a byproduct, the excited limbic system may create a brain status that is favorable for maintenance of REM sleep, leading to longer REM sleep duration.

One may wonder whether the offline performance gain by reward and sleep was due to merely the accumulation of reward provided during the acquisition stage. However, this possibility is unlikely because the threshold SOA at the initial training session was not significantly different between groups with or without reward in each experiment. The lack of effect of reward on the initial performance level was replicated in three independent experiments. Second, in Experiments 1 and 2, the increased offline performance gains were found only with the sleep groups, not with the wake groups. If reward does not interact with sleep, the effect of reward should have been significant not only for sleep groups but also for wake groups.

Is it possible that the effect of reward was merely unmasked visual adaptation that occurred during training session by sleep (9, 61)? This possibility is also unlikely. Visual adaptation is shown as decreased activity (62). If the observed effect of reward was due to merely a reduction in visual adaptation by sleep, given that visual adaptation is shown as decreased activity (62), the activation in the occipital regions during posttraining sleep should have been larger in the Reward group than in the No-reward group in Experiment 3. However, the results were opposite to this prediction. The results of Experiment 3 indicated that the overall effect of reward is inhibitory on visual processing. Moreover, the effect of reward was more apparent in the untrained region, not in the trained region, whereas visual adaptation emerged in the trained region (61). These findings suggest that the effect of reward on VPL is not due to merely the removal of visual adaptation by sleep but to a more active process involving the interaction between prefrontal reward processing and occipital visual processing. However, the interaction between reward and visual processing may be somehow suspended during wakefulness until posttraining sleep begins. Future research is needed to clarify what gates the interaction.

Was performance of the task affected by deprivation of water and food, not by reward per se? This possibility is unlikely for the following two reasons. First, if deprivation of water and food prior to the experiment had affected performance of the task, the initial performance level should have been different between groups with and without reward. However, as mentioned above, the threshold SOA in the initial training session was not significantly different across groups in any of the experiments. Second, if deprivation affected the task performance, water and food deprivation might have also affected subjective fatigue and sleepiness. However, the degree of subjective sleepiness was not significantly different between groups in either the presleep or postsleep test sessions (**Supplementary Table S3**). These data are inconsistent with the possibility that deprivation rather than reward affected performance.

Previous studies that investigated the relationship between reward values and selective remembering over sleep (59, 63, 64) have reported neither longer REM sleep nor increased activation during REM sleep. At least three possible distinctions might elucidate the difference between our study and previous studies. First, a different type of reward was used. While previous studies used money as a reward, the present study used water as a reward after fasting. Because a water reward after fasting could serve as a primary physical reinforcer, a water reward may have strongly impacted reward processing and/or the neural circuits involved in REM sleep. Second, we used larger differences in reward values across experimental conditions than in previous studies. In our study, one group was water-rewarded after fasting, while the other was not rewarded. By contrast, in most previous studies, a fine-graded difference in monetary reward values was used. For instance, a larger reward value was contrasted with a smaller reward value (59). The smaller difference in reward values might have obscured the impact on REM sleep. If true, this fact would further imply that the impact of reward during learning and memory on REM sleep may not be linear. Third, the tasks used were also different. Previous studies employed mostly associative memory tasks, whereas the present study employed VPL. Associative memories often involve hippocampal circuits (51, 52), whereas VPL of TDT used in the present study involves mostly the primary visual areas (25, 26, 30). The role of REM sleep might differ depending on the involved neural structures.

The present study used the TDT to investigate the interaction between the effects of reward and sleep on VPL. However, not all types of learning depend on the consolidation process during sleep (e.g., (65)). Additionally, whether reward during training is similarly effective on the consolidation of other types of VPL is unclear. More research is necessary to investigate whether the present findings can be generalized to other types of perceptual learning.

## MATERIALS AND METHODS

### Participants

A total of 69 young and healthy adults (47 (20 males and 27 females) in Experiment 1, 40 (16 males and 24 females) in Experiment 2, and 22 (10 males and 12 females) in Experiment 3) participated in the study. All participants had normal to corrected vision and were aged between 18 and 25 years old. Subjects gave their written informed consent for their participation after the purpose of the study was thoroughly described. The institutional review board of Brown University approved this study.

All participants were given a screening document to identify individuals who could safely refrain from eating or drinking 5 h prior to the experiment. None of the participants had prior experience with the task used in this study. Frequent video game players were excluded because prior research suggested that frequent gaming influences performance in VPL tasks (66–68). In addition, participants were required to have a regular sleep schedule, and anyone with a physical or psychiatric disease, who was currently under medication, or who was suspected to have a sleep disorder was excluded (16, 69, 70). Subjects were instructed to maintain regular sleep habits prior to the experiments and were asked to abstain from caffeine consumption during the day of the experiment. All participants were instructed that they could have water and food during the interval between sessions. Additional screening tests were conducted for polysomnographic experiments in Experiment 3 (see below).

### Design of Experiment 1

Due to an unexpected period of daytime napping, which could confound the results, one subject had to be excluded from the Wake & Reward group, leaving a total of 47 participants. The number of participants in each group was 12, except for the Wake & Reward group, which included 11 participants.

Subjects were trained and tested with a modified version of TDT (24, 71). Each trial began with a fixation point at the center of the display (1000 ms), which was followed by a briefly presented textured display (13 ms). After the disappearance of the textured display, a black screen was presented for a varying duration, followed by a mask stimulus for 100 ms. Mask stimuli were composed of randomly rotated V-shaped patterns. The blank interval between the onset of the textured stimulus and the mask is referred to as the stimulus-to-mask onset asynchrony (SOA), which varied from trial to trial. The textured display contained a central letter and a target triplet that consisted of 3 diagonal lines in the lower-left visual field quadrant in the background of horizontal lines. Participants were asked to report whether the central letter was ‘L’ or ‘T’ to ensure participants’ eye fixation for the letter task and then whether the target triplet was horizontally or vertically structured for the orientation task.

After the participant’s response, auditory feedback was provided for both the letter task and the orientation task in all groups. Furthermore, in the two reward groups (Sleep & Reward and Wake & Reward), participants received a droplet of water along with the auditory feedback provided for the orientation task for a correct response. Water was delivered to participants through a tube held in their mouth for the duration of the experiment (72). The water reward was provided during both the 1st and 2nd training sessions to control for the testing condition. Participants in the No-reward groups (Sleep & No-reward and Wake & No-reward) were not equipped with the feeding tube nor were they given water upon correct response for the orientation task.

The interval between the first and the second training sessions for each group was 12 h. Each training session consisted of 16 blocks, each of which had 39 trials with a constant SOA based on our previous study (67). Eight SOAs were considered: 400 ms, 180 ms, 160 ms, 140 ms, 120 ms, 100 ms, 80 ms and 60 ms. Each SOA was assigned 2 blocks (78 trials) (67). The total number of trials was 624, and approximately 1 h was required to complete each session.

The TDT stimulus subtended a 14°-by-14° visual angle. The position of each line segment in the background display jittered by 0–0.05° between trials. The stimulus was composed of 0.43°-by-0.07° (32 cd/m^2^) gray lines that were presented against a black background (0.5 cd/m^2^). The position of the orientation target varied slightly between each trial but was consistently presented within the lower-left quadrant within a 3°–5° visual angle from the center of the display.

The stimuli were presented using Psychophysics Toolbox (73, 74) for MATLAB® (The MathWorks, Natick, MA) on a Macintosh G5 computer. The stimuli appeared on a 19″ CRT monitor with a resolution of 1024 by 768 pixels at a refresh rate of 85 Hz. Participants’ heads were restrained with a chin rest, and the viewing distance was 57 cm. All experiments were conducted in a dimly lit room.

For each session, we obtained the 75% threshold SOA as a behavioral measure. First, we obtained the correct response ratio for the peripheral orientation task computed for each SOA and then fitted the data to a logistic psychometric function (75). The performance improvement% was defined as follows: (threshold SOA of the training session - threshold SOA of the test session) / threshold SOA of the training session.

### Design of Experiment 2

The total sample for the study was 40 participants. The number of participants was 11 for the Sleep & Reward, Sleep & No-reward, and Wake & No-reward groups and 7 for the Wake & Reward group.

Subjects were trained and tested with the same version of the TDT as in Experiment 1 (24, 71). The training session consisted of 16 blocks, each of which had 39 TDT trials, as in Experiment 1. The range of SOAs used was also the same as in Experiment 1. In the Reward groups only, participants were asked to refrain from eating or drinking for 4-5 h prior to the training session. Participants were provided a water reward upon correct response for the peripheral orientation task with the same method and apparatus as used in Experiment 1. Both test sessions T1 and T2 included 16 blocks, each of which consisted of 5 trials so that no improvement should occur during the test sessions. Thus, each test session consisted of 80 trials in total (10 trials per SOA), which took approximately 10 min. Water reward was given only during the training session. No water reward was given during the T1 and T2 test sessions so that both the Reward and No-reward groups had the same procedure during the test sessions.

The performance improvement over the wake/sleep period was calculated based on the threshold SOA in test sessions T1 and T2. First, for each test session, the 75% threshold SOA was obtained. Second, the threshold SOA of session T2 was subtracted from the threshold SOA of session T1, then divided by the threshold SOA of session T1 session and multiplied by 100.

### Design of Experiment 3

Because Experiment 3 was designed to examine brain activation during sleep, we applied screening tests for the participants in addition to those conducted in Experiment 1 to select subjects whose sleep habits were more consistent. Participants were required to have a regular sleep schedule of approximately 7-8 h of sleep with a bedtime of 11 pm to 12 am. We conducted a questionnaire to ask whether a subject may have symptoms related to sleep complaints or sleep disorders, including insomnia and narcolepsy. None of the subjects reported sleep complaints or disorders. In addition, the Pittsburg Sleep Quality Index (PSQI) (33) and the Munich Chronotype Questionnaire (MCTQ) (34, 35) were used to characterize subjects’ habitual sleep quality and individual chronotypes.

Participants were asked to arrive at the laboratory at noon. After a 1-h period of TDT training and the first 10-min test session T1, participants were allowed to rest for a period of approximately 30 min while electrodes were applied for PSG. A 120-min nap began at approximately 2 pm for all participants. Test session T2 was conducted at approximately 4:30 pm, once participants were awake and unequipped with the PSG electrodes.

All the procedures and parameters for the training and test sessions were the same as those in Experiment 2. A water reward was given only during the training session, and no water reward was given during test sessions T1 and T2, so both the Reward and No-reward groups had the same procedure during the test sessions.

The performance improvement over the sleep period was calculated based on the threshold SOA in test sessions T1 and T2. First, for each test session, the 75% threshold SOA was obtained. Second, the threshold SOA of session T2 was subtracted from the threshold SOA of session T1 session, then divided by the threshold SOA of session T1 session and multiplied by 100.

Subjective sleepiness was measured using SSS at the beginning of test sessions T1 and T2 (29).

One may wonder whether the sleep structures, including REM phases, were comparable between the groups before training. To confirm that there were no significant differences between the Reward and No-reward groups in terms of basic sleep structures, including the REM phase, in Experiment 3, we conducted additional analyses on the questionnaires and the data from the sleep sessions. First, we analyzed the PSQI (33) and the MCTQ (34, 35). No significant differences were observed between groups in terms of any of the habitual sleep parameters (**Supplementary Table S7**). To further confirm that the phases of REM sleep were comparable across groups, we measured the latency to REM sleep from the posttraining sleep session (Table 1). First, no subject showed sleep-onset REM sleep. Second, the latency to REM sleep was nonsignificant between groups (unpaired t-test, t(18)=0.84, p=0.414). These results indicate that the REM phases were comparable between groups.

### PSG in Experiment 3

Subjects slept in a soundproof and shielded room monitored by PSG, which consisted of electroencephalogram (EEG), electrooculogram (EOG), and electromyogram (EMG). EEG was recorded at 64 scalp sites according to the 10% electrode position (76) using active electrodes (actiCap, Brain Products, LLC) with a standard amplifier (BrainAmp Standard, Brain Products, LLC). The online reference was Fz, which was rereferenced to the average of the left and right mastoids offline after the recording for postprocessing. The sampling frequency was 500 Hz, and the impedance was kept below 20 kΩ. The active electrodes included a new type of integrated impedance converter, which allowed them to transmit the EEG signal with significantly lower levels of noise than traditional passive electrode systems. The data quality with active electrodes was as good as 5 kΩ using passive electrodes, which were used for EOG and EMG (BrainAmp ExG, Brain Products, LLC). Horizontal EOG was recorded from 2 electrodes placed at the outer canthi of both eyes. Vertical EOG was measured from 4 electrodes placed 3 cm above and below both eyes. EMG was recorded from the mentum or chin. The impedance was kept below 10 kΩ for the passive electrodes. Brain Vision Recorder software (Brain Products, LLC) was used for PSG recording. The data were filtered between 0.1 and 40 Hz, and a 30 Hz high-pass filter was applied to the data before scoring to reduce noise.

Sleep stages were scored for every 30-s epoch according to the standard criteria (77, 78) into wakefulness (Wake), NREM stage 1 sleep (N1), NREM stage 2 sleep (N2), NREM stage 3 sleep (N3), and REM stage sleep. Sleep-onset was defined as the first appearance of N2, in accordance with prior work (69, 70).

EEG was fast Fourier transformed in 5-s epochs and smoothed with a tapered cosine window throughout each of N2, N3, and REM sleep stages. Six epochs were used to yield the averaged spectral power (µV^2^) of 30 s for each channel for each of the following frequency bands: delta (1-4 Hz), theta (5-9 Hz), alpha (10-12 Hz) and sigma (13-16 Hz).

To normalize individual differences in the absolute power (79–81), we obtained the power per EEG channel for each frequency band per 30 s for each of the NREM and REM sleep stages. Second, the average power across all EEG channels for each frequency band was computed for the NREM (N2 and N3) and REM sleep stages. Third, for each EEG channel, the relative power density (%) was obtained by dividing each absolute power by the average power. For instance, if a given EEG channel’s power was 120 µV^2^ and the average power across all channels was 100 µV^2^, then the power density for that channel would be 120%. Thus, the power density is normalized across all channels for each frequency band by NREM or REM sleep stage.

We preselected prefrontal (F1, F2, AF3, AF4) and occipital (PO3, PO4, PO7, PO8) EEG channels for subsequent analyses. Prefrontal EEG channels have been reported to be sensitive to reward processing in the dorsolateral and ventromedial prefrontal regions in a previous study (23), in which dedicated source localization was conducted in combination with EEG and magnetic resonance imaging. In addition, because the present task is known to show trained-location specificity in learning (24, 71, 75) and the target orientation was always presented in the left lower visual field quadrant, the PO4 and PO8 channels in the right occipital region were assigned to cover the trained location, and the PO3 and PO7 channels in the left occipital region were assigned to cover the untrained location, based on a previous study (8). The EEG data during REM sleep of two subjects in the No-reward group were too noisy; thus, the number of subjects for the No-reward group was 9, whereas it was 11 for the Reward group when we analyzed spontaneous oscillations during REM sleep. No data were omitted for the analyses on spontaneous oscillations during NREM sleep.

## ACKNOWLEDGMENTS

This work was supported by NIH (R21EY028329, R01EY019466, R01EY027841, T32EY018080) and BSF2016058.

