## Supplementary file for "Reward does not facilitate visual perceptual learning until sleep occurs"

**This PDF file includes:**

Figures S1 to S4
Tables S1 to S7
SI References

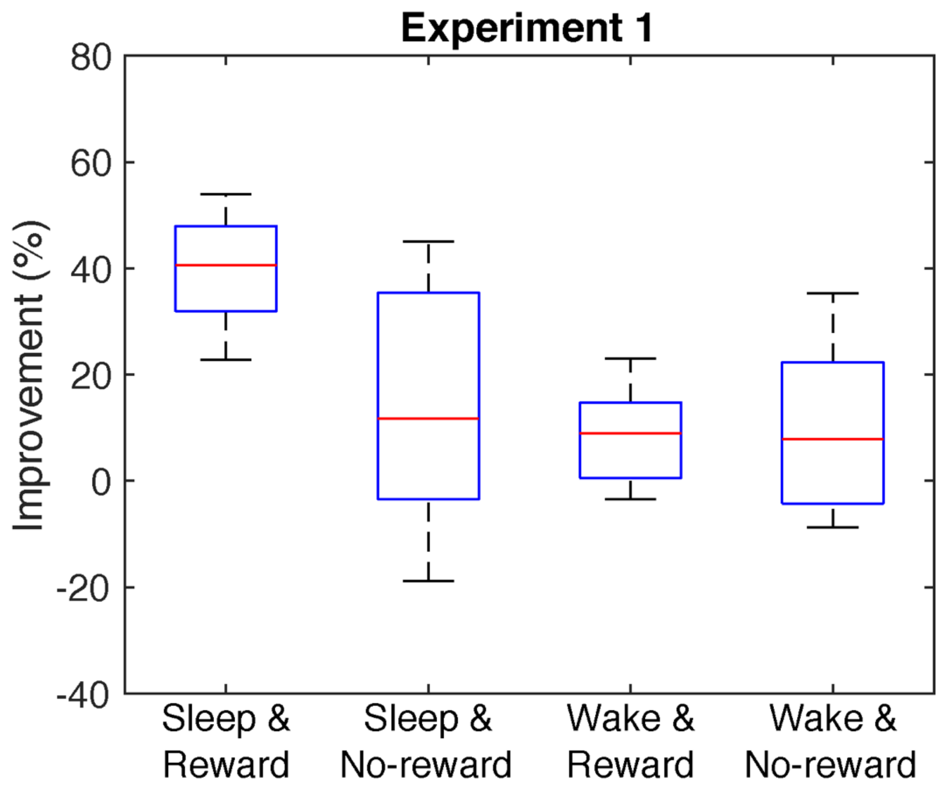

**Fig. S1.** Performance improvement for Experiment 1. The bottoms and tops of the blue boxes show the 25th and 75th percentiles (the lower and upper quartiles), respectively; the inner red band shows the median; the whiskers show the maximum and minimum of the data that are not considered to be outliers based on Grubbs' test (Alpha=.05, two-sided).

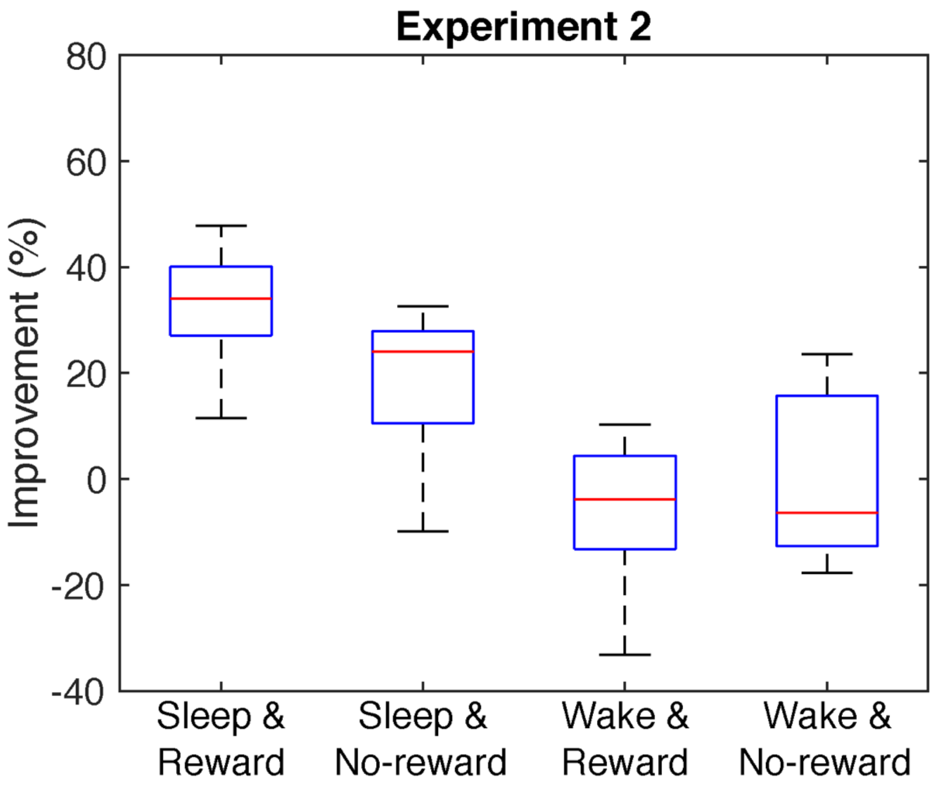

**Fig. S2.** Performance improvement for Experiment 2. The bottoms and tops of the blue boxes show the 25th and 75th percentiles (the lower and upper quartiles), respectively; the inner red band shows the median; the whiskers show the maximum and minimum of the data that are not considered to be outliers based on Grubbs' test (Alpha=.05, two-sided).

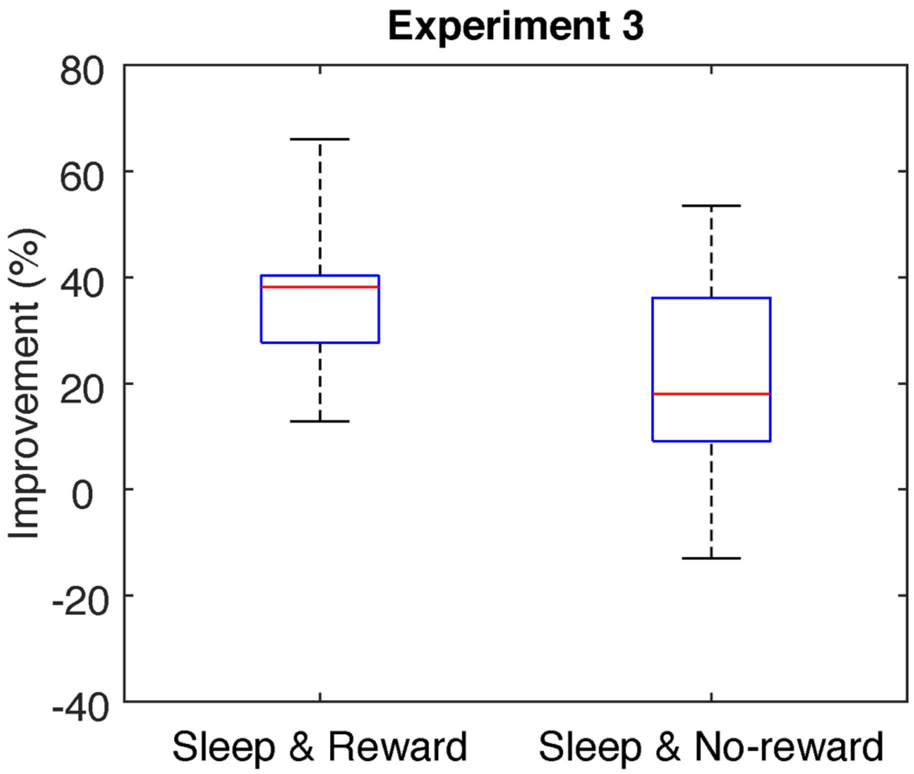

**Fig. S3.** Performance improvement for Experiment 3. The bottoms and tops of the blue boxes show the 25th and 75th percentiles (the lower and upper quartiles), respectively; the inner red band shows the median; the whiskers show the maximum and minimum of the data that are not considered to be outliers based on Grubbs' test (Alpha=.05, two-sided).

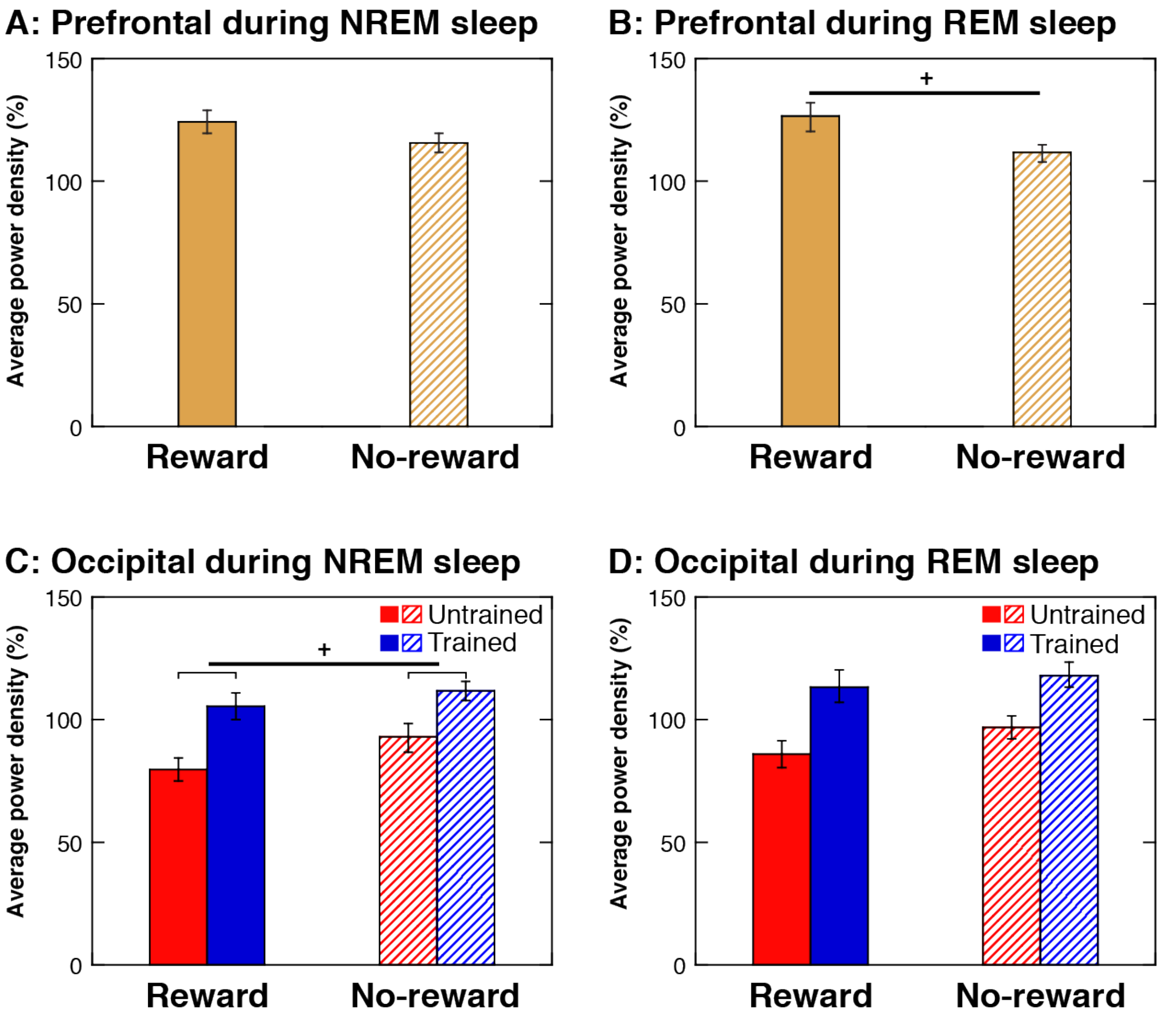

**Fig. S4.** Spontaneous brain activities for each brain region averaged across the 4 frequency bands. The effect of reward on the averaged power densities (±SEM) at the prefrontal (A and B) and occipital (C and D) regions during NREM sleep and REM sleep. Filled bars represent the Reward group, and hatched bars represent the No-reward group. Yellow bars denote the prefrontal (A and B), red bars denote the untrained occipital (C and D) and blue bars denote the trained occipital regions (C and D). A: The power density in the prefrontal region during NREM sleep. B: The power density in the prefrontal region during REM sleep. C: The power density in the occipital region during NREM sleep. D: The power density in the occipital region during REM sleep.

**Table S1.** Threshold SOA (ms) for Experiment 1

| | 1st training | 2nd training |
|---|---|---|
| Sleep & Reward | 156.7 ± 7.58 | 92.3 ± 5.52 |
| Sleep & No-reward | 140.0 ± 7.91 | 120.4 ± 7.77 |
| Wake & Reward | 144.1 ± 6.83 | 131.1 ± 7.06 |
| Wake & No-reward | 145.7 ± 7.70 | 133.8 ± 8.76 |

Values are the mean ± SE.

**Table S2.** Threshold SOA (ms) for Experiment 2

| | Training | T1 | T2 |
|---|---|---|---|
| Sleep & Reward | 146.6 ± 14.42 | 148.6 ± 16.67 | 114.5 ± 14.46 |
| Sleep & No-reward | 137.1 ± 13.06 | 135.5 ± 12.33 | 115.5 ± 10.39 |
| Wake & Reward | 147.3 ± 13.14 | 130.0 ± 11.68 | 145.5 ± 15.10 |
| Wake & No-reward | 142.5 ± 12.58 | 142.9 ± 10.92 | 144.6 ± 11.18 |

Values are the mean ± SE.

**Table S3.** Sleepiness for Experiment 3

| | T1 | | | T2 | | |
|---|---|---|---|---|---|---|
| Reward | 1.45 | ± | 0.16 | 1.91 | ± | 0.21 |
| No-reward | 1.55 | ± | 0.21 | 2.00 | ± | 0.23 |

Values are the mean ± SE.

**Table S4.** Threshold SOA (ms) for Experiment 3

| | Training | T1 | T2 |
|---|---|---|---|
| Reward | 141.8 ± 11.29 | 144.4 ± 17.43 | 86.5 ± 9.72 |
| No-reward | 137.0 ± 7.52 | 138.0 ± 12.07 | 106.9 ± 11.64 |

Values are the mean ± SE.

**Table S5.** 4-way ANOVA and post hoc results in Experiment 3. We highlighted the significant results (p<0.05).

### A. Interactions and main effects

| | | |
|---|---|---|
| **4-way interaction** | F(6,108)=3.234 | p=0.006 |
| Stage * Freq * REGION * Group | | |
| **3-way interaction** | | |
| Stage * Freq * REGION | F(6,108)=3.818 | p=0.002 |
| Freq * REGION * Group | F(6,108)=1.78 | p=0.11 |
| Stage * REGION * Group | F(2,36)=0.191 | p=0.827 |
| Stage * Freq * Group | F(3,54)=2.255 | p=0.092 |
| a main effect | | |
| Stage | F(1,18)=6.73 | p=0.018 |
| Freq | F(3,54)=3.663 | p=0.018 |
| REGION | F(2,36)=20.608 | p=0.000 |
| Group | F(1,18)=0.229 | p=0.638 |
| 2-way interaction | | |
| Stage * Group | F(1,18)=1.35 | p=0.261 |
| Freq * Group | F(3,54)=0.683 | p=0.566 |
| REGION * Group | F(2,36)=3.034 | p=0.061 |
| Stage * Freq | F(3,54)=11.651 | p=0.000 |
| Stage * REGION | F(2,36)=2.491 | p=0.097 |
| Freq * REGION | F(6,108)=9.001 | p=0.000 |

### B. Group (reward) effect

| Stage | Freq | REGION | df | F | Sig. |
|---|---|---|---|---|---|
| NREM | Delta | Frontal | F(1, 18) | 0.139 | 0.713 |
| NREM | Delta | Untrained | F(1, 18) | 0.742 | 0.400 |
| NREM | Delta | Trained | F(1, 18) | 1.433 | 0.247 |
| NREM | Theta | Frontal | F(1, 18) | 2.049 | 0.169 |
| NREM | Theta | Untrained | F(1, 18) | 1.230 | 0.282 |
| NREM | Theta | Trained | F(1, 18) | 0.029 | 0.866 |
| NREM | Alpha | Frontal | F(1, 18) | 1.931 | 0.182 |
| NREM | Alpha | Untrained | F(1, 18) | 0.616 | 0.443 |
| NREM | Alpha | Trained | F(1, 18) | 0.910 | 0.353 |
| NREM | Sigma | Frontal | F(1, 18) | 3.893 | 0.064 |
| NREM | Sigma | Untrained | F(1, 18) | 12.434 | 0.002 |
| NREM | Sigma | Trained | F(1, 18) | 1.442 | 0.245 |
| REM | Delta | Frontal | F(1, 18) | 1.684 | 0.211 |
| REM | Delta | Untrained | F(1, 18) | 3.750 | 0.069 |
| REM | Delta | Trained | F(1, 18) | 1.533 | 0.232 |
| REM | Theta | Frontal | F(1, 18) | 5.423 | 0.032 |
| REM | Theta | Untrained | F(1, 18) | 4.822 | 0.041 |
| REM | Theta | Trained | F(1, 18) | 0.236 | 0.633 |
| REM | Alpha | Frontal | F(1, 18) | 5.256 | 0.034 |
| REM | Alpha | Untrained | F(1, 18) | 1.051 | 0.319 |
| REM | Alpha | Trained | F(1, 18) | 2.340 | 0.144 |
| REM | Sigma | Frontal | F(1, 18) | 4.718 | 0.043 |
| REM | Sigma | Untrained | F(1, 18) | 0.188 | 0.670 |
| REM | Sigma | Trained | F(1, 18) | 1.650 | 0.215 |

**Table S6.** Correlation coefficients and p-values between sleep parameters and the threshold SOA

| | Training | | T1 | |
|---|---|---|---|---|
| | *r* | *p* | *r* | *P* |
| Stage W | 0.06 | 0.80 | -0.01 | 0.97 |
| Stage N1 | -0.23 | 0.30 | -0.34 | 0.12 |
| Stage N2 | -0.03 | 0.89 | -0.01 | 0.99 |
| Stage N3 | 0.17 | 0.45 | 0.18 | 0.42 |
| Stage REM | 0.15 | 0.51 | -0.02 | 0.93 |
| SOL | -0.15 | 0.50 | -0.12 | 0.59 |

SOL, sleep-onset latency.

**Table S7.** Habitual sleep parameters for Experiment 3

| | Reward | | | No-reward | | |
|---|---|---|---|---|---|---|
| Bed time[a] | 23:35 | ± | 0:09 | 23:25 | ± | 0:10 |
| Wake-up time[b] | 7:16 | ± | 0:10 | 7:09 | ± | 0:12 |
| Midsleep time[c] | 3:51 | ± | 0:10 | 3:50 | ± | 0:07 |
| Habitual sleep efficiency (%)[d] | 95.9 | ± | 0.89 | 97.1 | ± | 1.10 |

Values are the mean ± SE.

**a & b**, Habitual bed time and wake-up time were obtained from the PSQI (1). There was no significant difference between groups in terms of bed time ($t(20)=0.62$, $p=0.510$) or wake-up time ($t(20)=0.43$, $p=0.673$).

**c**, The 'midsleep' time, which is the midpoint between sleep-onset and wake-up time, was obtained from the Munich Chronotype Questionnaire (MCTQ; (2, 3)). The midsleep time is significantly correlated with the circadian phase sleep (3, 4) that influences the timing of REM sleep. Therefore, a significant difference in midsleep times between the groups would suggest that the phases of the occurrence of REM sleep were dramatically different between the groups before training. However, no significant difference between groups was observed ($t(20)=0.07$, $p=0.949$).

**d**, The habitual sleep efficiency (%) was measured as [(Number of hours slept / Number of hours spent in bed) X 100)] for each group using the PSQI (1). No significant difference in sleep efficiency was observed between groups ($t(20)=0.838$, $p=0.412$).
